# Local association of *Trypanosoma cruzi* chronic infection foci and enteric neuropathic lesions at the tissue micro-domain scale

**DOI:** 10.1101/2021.03.09.434577

**Authors:** Archie A. Khan, Harry C. Langston, Fernanda C. Costa, Francisco Olmo, Martin C. Taylor, Conor J. McCann, John M. Kelly, Michael D. Lewis

## Abstract

Gastrointestinal (GI) disease affects a substantial subset of chronic Chagas disease (CD) patients, but the mechanism of pathogenesis is poorly understood. The lack of a robust, predictive animal model of chronic *T. cruzi* infection that exhibits functional digestive disease has held back research. To address this, we combined GI tracer assays and bioluminescence *in vivo* infection imaging systems for diverse parasite strains to discover models exhibiting chronic digestive transit dysfunction. We identified the colon as a specific site of both tissue parasite persistence and delayed transit. Digestive CD mice exhibited significant retention of faeces in both sated and fasted conditions. Histological and immunofluorescence analysis of the enteric nervous system (ENS) revealed a dramatic reduction in the number of neurons and a loss of immunoreactivity of the enteric neural network in the colon. This model therefore recapitulates key clinical manifestations of human digestive CD. We also exploited dual bioluminescent-fluorescent parasites to analyse rare chronic infection foci in the colon at the single cell level, revealing co-localisation with ENS lesions. This indicates that long-term *T. cruzi*-host interactions in the colon drive pathogenesis and thus chronic disease may be preventable using anti-parasitic chemotherapy.

## Introduction

Chagas disease (CD) is caused by infection with the protozoan parasite *Trypanosoma cruzi*, which affects approximately 6 million people. There are two principal forms of CD, cardiac and digestive. The most prevalent cardiac presentations include myocarditis, fibrosis, arrhythmias, microvascular abnormalities, progressive heart failure and sudden death ^1^. Cardiac CD has been the subject of intensive experimental research and many predictive animal models are available to support translation into the clinic. Human digestive CD (DCD) is characterised by progressive dilatation and dysfunction of sections of the GI tract ^2,3^. Symptoms include achalasia, abdominal pain, constipation and faecaloma. Eventually, massive organ dilatation results in megasyndromes, usually of the colon or oesophagus. Dilatation is associated with loss of enteric neurons leading to peristaltic paralysis and smooth muscle hypertrophy. Treatments are limited to dietary and surgical interventions. The lack of a robust small animal model of enteric CD has been a major block on basic and translational research.

Of symptomatic CD patients, ~65% have cardiomyopathy, 30% enteropathy and 5% have both, with digestive disease most common in Bolivia, Chile, Argentina and Brazil ^1^. Anti-parasitic chemotherapy has not been considered justifiable for *T. cruzi-*positive individuals with digestive symptoms but normal heart function ^4^. The primary reason is that no clinical trials have addressed treatment efficacy in the context of digestive outcomes, and there are little to no experimental data on which to base such trials. Molecular and cellular explanations of DCD pathogenesis also lag far behind the advances made for Chagas cardiomyopathy.

The lack of progress in developing treatments for DCD is also connected to the prevailing view that megasyndromes result from irreversible enteric denervation, specifically during the *acute* phase of infection ^5,6^, in which anti-parasitic inflammatory responses are thought to cause iNOS-dependent collateral damage to neurons, leading to aganglionosis ^5,7^. Further age-related nerve degeneration is posited to gradually unmask these parasite-driven losses, leading to progressive organ dysfunction on a timescale of years to decades ^5^. Recently, several lines of evidence have converged to question this theory. Experimental infection imaging studies in mice revealed that GI tract is in fact a long-term reservoir of *T. cruzi* infection ^8,9^. Adult enteric neurogenesis has been described in response to chemically-mediated tissue injury ^10^ and in the steady state ^11^. A series of advances has also highlighted previously unappreciated levels of interconnectedness between the gut’s immune and nervous systems ^12–14^. We therefore hypothesized that host-parasite interactions in the chronically infected gut might impact continuously on the enteric nervous system (ENS) and musculature to drive DCD pathogenesis.

Here, we screened a series of parasite and mouse strain combinations to identify several models with significant digestive motility dysfunction. Using a combination of bioluminescence and fluorescence *in vivo* and *ex vivo* imaging techniques we demonstrate that chronic *T. cruzi* persistence, gut motility delay and enteric neuronal damage are co-localised within discrete foci in the colonic muscularis. This indicates that DCD tissue pathology and transit dysfunction are likely driven by *T. cruzi* persistence in the colon and the associated chronic inflammatory response. DCD should therefore be considered as potentially preventable by anti-parasitic chemotherapy. It also opens the way to investigate the molecular and cellular basis of pathogenesis and *T. cruzi* immune evasion.

## Results

### Screening *T. cruzi* mouse infection models for delayed digestive transit times

We previously developed a series of mouse models of *T. cruzi* infection based on parasites transgenically expressing the luciferase variant *Ppy*RE9h, which serves as an orange-red light emitting *in vivo* reporter protein ^15^. Host-parasite combinations of BALB/c and C3H/HeN mice and TcVI-CL Brener (CLBR) and TcI-JR strain parasites permit long-term tracking of the course and distribution of infections in individual animals (Figure 1a). These models, which exhibit a spectrum of Chagas heart disease severities ^9^, were selected to screen for gastrointestinal (GI) transit time delays, a common feature of DCD, by oral feeding of a red dye tracer (carmine). Three of the four host-parasite combinations took significantly longer than control uninfected mice to pass the tracer at acute phase, 3 weeks post-infection (p.i.), and/or at 6 weeks p.i. transition phase (Figure 1b). During the early chronic phase, 12 and 18 weeks p.i., only the TcI-JR-infected C3H mice displayed the delay phenotype, which became markedly more severe as the infection developed into the later chronic phase at 24 and 30 weeks p.i. Milder, but still significant transit delay phenotypes also emerged in the other three models.

**Figure 1:**
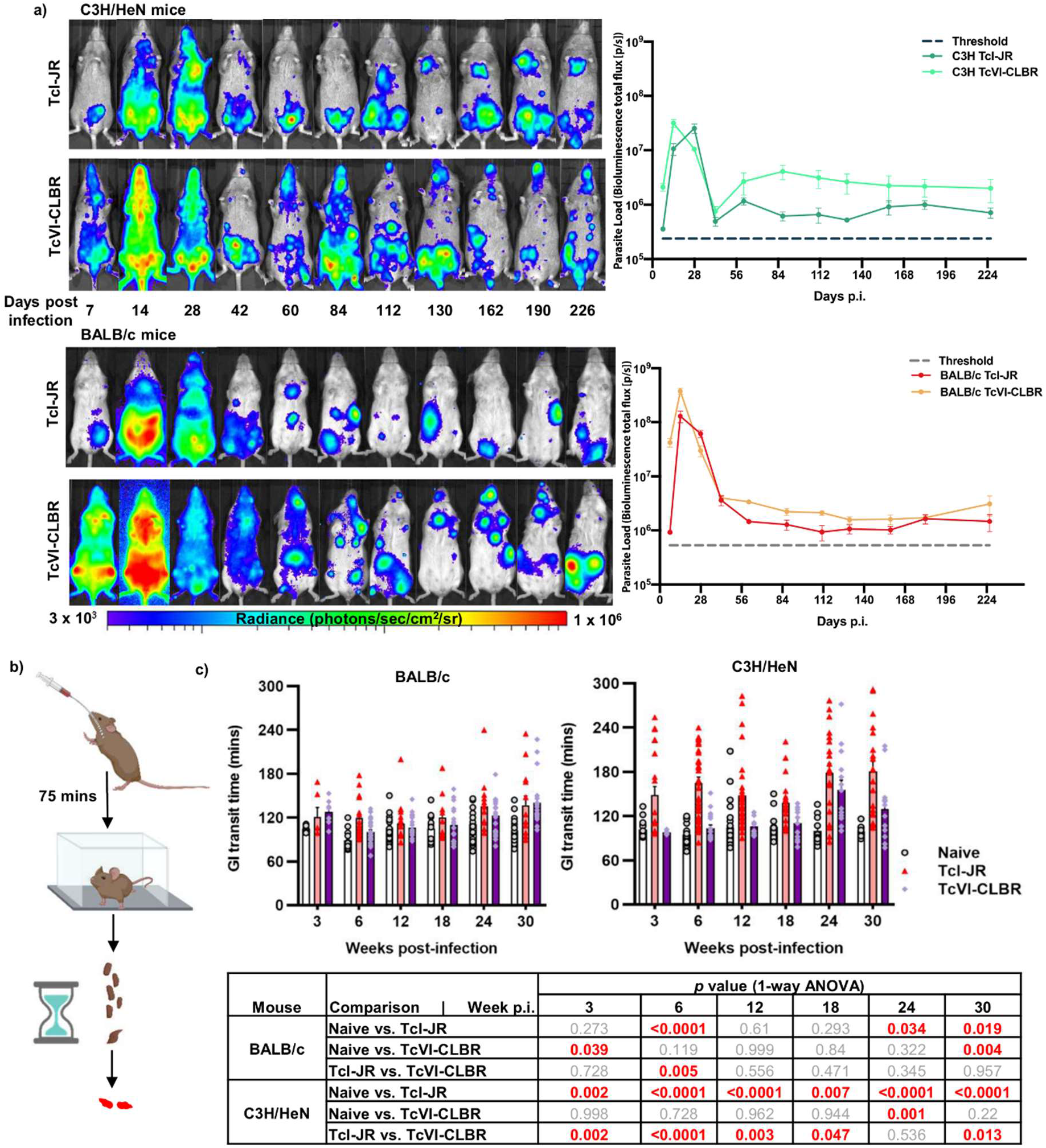
Bioluminescence imaging *T. cruzi* infection models and digestive transit dysfunction screen. **a)** Ventral images of female C3H/HeN (top panels) and BALB/c mice (bottom panels) representing TcI-JR (1^st^ and 3^rd^ panel) and TcVI-CLBR (2^nd^ and 4^th^ panel) course of infection. Images were captured using *in vivo* bioluminescence imaging. Overlaid log-scale pseudocolour heat maps are representative of bioluminescence intensity; the log-scale range is indicated in units of radiance. Adjacent line plots show parasite load represented in average bioluminescence of TcI-JR C3H/HeN (*n* = 10-24), TcVI-CLBR C3H/HeN (*n* = 5-12), TcI-JR BALB/c (*n* = 5-12) and TcVI-CLBR BALB/c (*n* = 9-22) infected mice against days post infection (p.i.). Limit of detection of bioluminescence is indicated as threshold by dashed line. **b)** Schematic diagram of the carmine red-dye assay to measure gastrointestinal (GI) transit time delay in mice. **c)** Bar plots show GI transit time vs. weeks post-infection (p.i.) of BALB/c (left) and C3H/HeN (right) mice in the following groups: naive control BALB/c (*n* = 8-18), TcI-JR BALB/c (*n* = 6-17), TcVI-CLBR BALB/c (*n* = 10-29), uninfected naive control C3H/HeN (*n* = 12-35), TcI-JR C3H/HeN (*n* = 18-38) and TcVI-CLBR C3H/HeN (*n* = 6-17). Table (bottom) summarises statistical comparisons of GI transit time delay between groups. All statistically significant values are highlighted (red). Data are expressed as mean ± SEM. Statistical significance was tested using one way ANOVA followed by Tukey’s HSD test.

*T. cruzi* as a species encompasses a high level of genetic diversity structured across six major lineages ^16–18^. To test whether and at what level the strong digestive transit delay phenotype in C3H mice was conserved, we tested a further nine *T. cruzi* strains from five lineages (4x TcI, 1x TcII, 1x TcIII, 1x TcIV and 1x TcVI) using the carmine transit assay (Supplementary Figure 1). Two more strains were identified showing evidence of delayed transit: TcI-SN3 and TcVI-Peru. This type of pathology is therefore a relatively rare, strain-specific trait in *T. cruzi*. It occurs in both TcI and TcVI strains, but is not conserved within lineages.

### Parasite persistence within the GI tract

We selected the TcI-JR-infected C3H mouse as the most suitable model of experimental chronic DCD. The transit time delay in these animals (Figure 1b) did not show a correlation with the overall parasite burden, which dropped by approximately two orders of magnitude from the acute peak to the level seen in the chronic phase (Figure 1a). Much of the bioluminescence signal in whole animal imaging derives from parasites in the skin ^8,9^, so we quantified organ-specific parasite loads using *ex vivo* imaging at 3, 6 and 30 weeks p.i. (Figure 2a). Parasitism was consistently detected in the GI tract, in foci distributed from the stomach to the rectum, being relatively more intense in the stomach and large intestine compared to the small intestine (Figure 2a, 2b). All sites exhibited significantly lower parasite loads in the chronic than acute phase (Figure 2b). Thus, GI transit delays were associated with local persistence of *T. cruzi.* There was a positive correlation between endpoint GI parasite loads and the severity of transit delay during the acute phase (3 weeks p.i.), but there were no such quantitative associations in the transition (6 weeks) or chronic (30 weeks) phases (Figure 2c).

**Figure 2:**
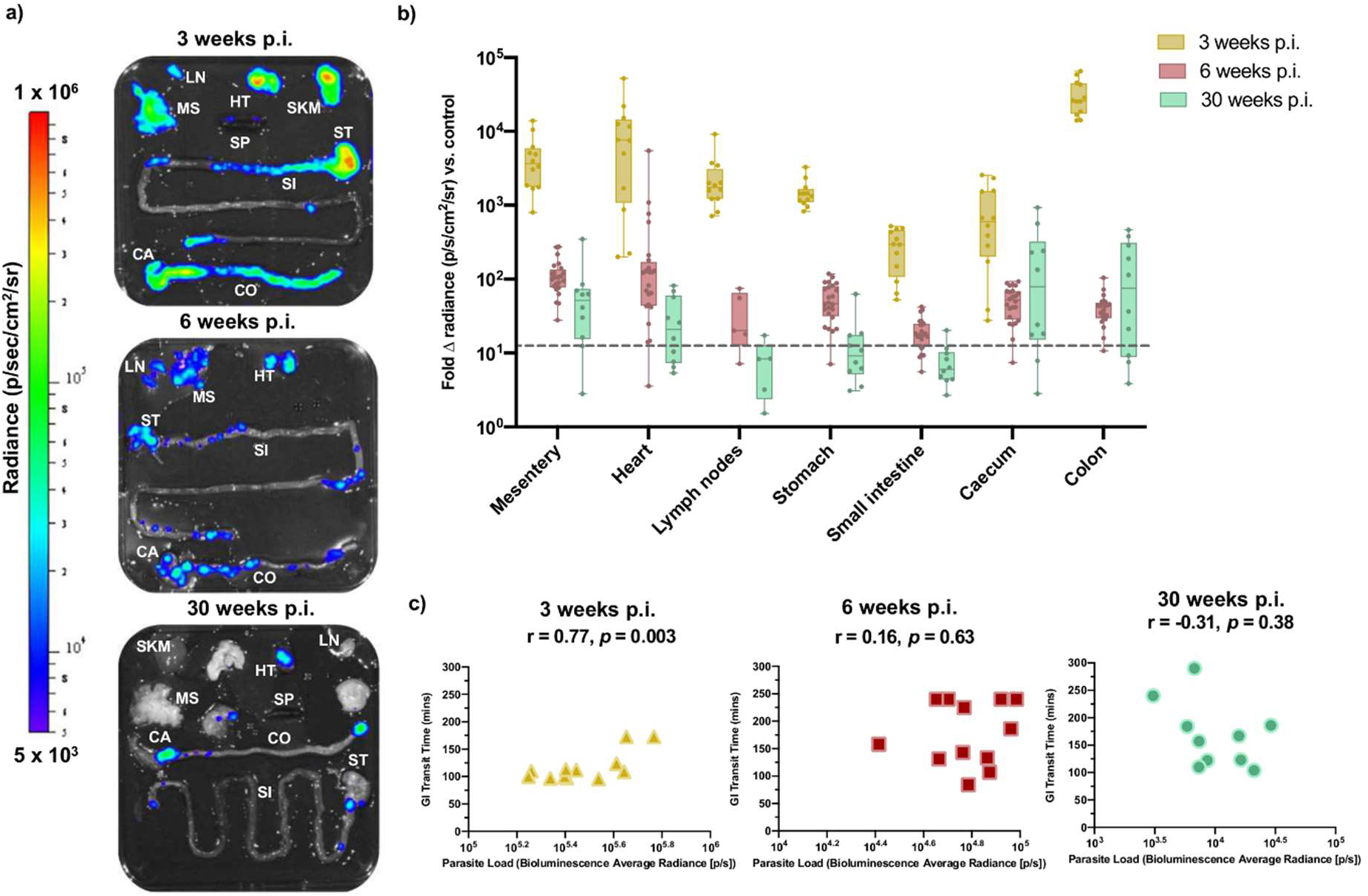
Tissue parasite distribution kinetics in TcI-JR-infected C3H/HeN mice. **a)** Representative images show parasite distribution in different organ tissue (lymph nodes-LN, gut mesenteric tissue-MS, heart-HT, spleen-SP, skeletal muscle-SKM, stomach-ST, small intestine-SI, caecum-CA and colon-CO) of a TcI-JR infected C3H mouse at 3, 6 and 30 weeks post-infection (p.i.) using *ex-vivo* bioluminescence imaging. Overlaid log-scale pseudocolour heat maps are representative of bioluminescence intensity; the log-scale range is indicated in units of radiance. **b)** Box-plots show infection intensity of different organ tissue at 3 (*n* = 12 per group), 6 (*n* = 24 per group; *n* = 5 lymph nodes) and 30 (*n* = 10 per group; *n* = 5 lymph nodes) weeks p.i. Data points are expressed as fold change in bioluminescence vs. naïve controls. Limit of detection is denoted as dashed line. The horizontal line within each box indicates median and the whiskers denotes minimum and maximum values of each dataset. **c)** Scatter plots show correlation between gastrointestinal transit time and end-point parasite densities expressed as bioluminescence in radiance at 3 (*n* = 10), 6 (*n* = 12) and 30 (*n* = 10) weeks p.i.; r denotes Pearson’s correlation coefficient and p-value represents a measure of statistical significance.

### Regional dissection of the transit delay phenotype reveals localisation to the colon

The transit time delay seen in infected animals was not explained by differences in body weight or intestine length (Supplementary Figure 2). This suggested a functional impairment to peristalsis, as seen in human DCD. Our next aim was to determine the digestive tract region(s) in which the transit time delay was localised. To do this we fed mice with red and green fluorescent tracers (Rhodamine dextran and 70kDa FITC dextran, respectively) at variable time intervals prior to *ex vivo* imaging. An interval of 5 minutes was used to test whether stomach emptying was delayed. No significant differences were detected in infected animals compared to controls (Figure 3a), either at 6 or 30 weeks p.i. There was a significant difference in stomach weight at 6 weeks p.i. (Figure 3b), which may indicate increased retention of matter more solid than the tracer.

**Figure 3:**
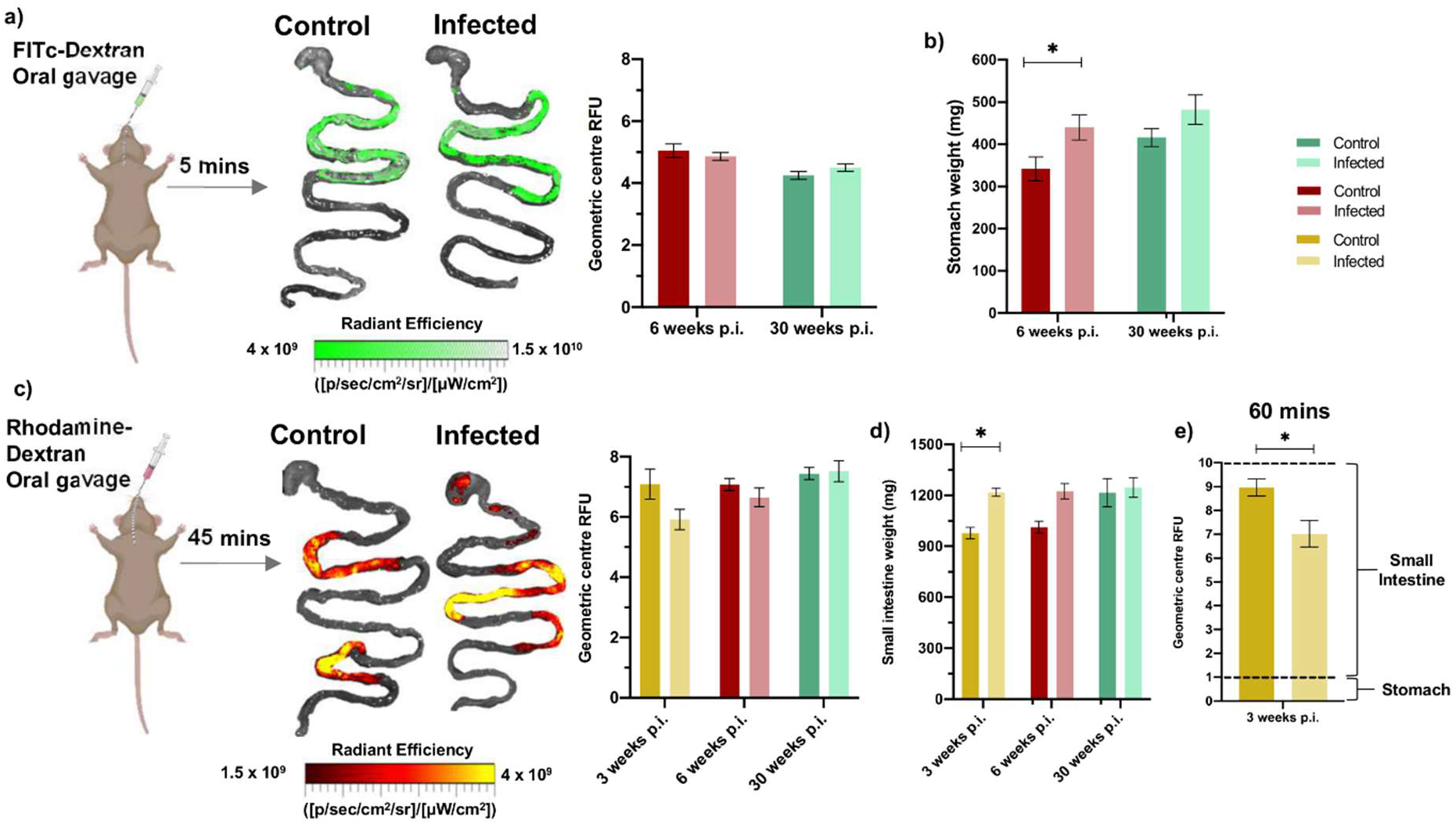
Fluorescent tracer imaging assays for stomach emptying and small intestine transit. **a)** Schematic diagram of a mouse receiving oral gavage of a green fluorescent marker, 70-kDa FITc-dextran, 5 minutes prior to termination to trace stomach emptying delay during infection. Representative images of stomach and small intestine are superimposed with traces of 70-kDa FITc-dextran travelling through stomach into small intestine to show transit difference between control and TcI-JR C3H/HeN infected mice. Linear-scale pseudocolour heat map shows minimum and maximum fluorescence intensity of 70-kDa FITc-dextran. Quantification of FITc-dextran fluorescence in control naïve and TcI-JR C3H/HeN is shown in the adjacent bar plot at 6 (*n* = 12 per group) and 30 (*n* = 5 per group) weeks post-infection (p.i.). Fluorescence is expressed as geometric centre which is centre mass of the marker. **b)** Bar plot shows post-mortem weights of stomach with contents at 6 (*n* = 7 per group) and 30 (*n* = 5 per group) weeks p.i. **c)** Similar schematic diagram and bar plot at 3 (*n* = 4 per group), 6 (*n* = 4 per group) and 30 (*n* = 5 per group) weeks p.i. using a red fluorescent marker, rhodamine-dextran, to target small intestine transit. Linear-scale pseudocolor heat map shows minimum and maximum fluorescence intensity of rhodamine-dextran. **d)** Small intestine weights shown in bar plot at 3 (*n* = 4 per group), 6 (*n* = 7 per group) and 30 (*n* = 5 per group) weeks p.i. **e)** Bar plot shows quantification of rhodamine-dextran fluorescence administered 60 minutes before termination of mice at 3 weeks p.i. (*n* = 4 per group). Dashed lines on bar plots show the GI segment number corresponding to the geometric centre score (0-1= stomach, 1-10 = small intestine). Data are expressed as mean ± SEM. Statistical significance was tested using unpaired two-tailed Student’s t test (\**P* < 0.05).

To measure small intestine dysfunction, we initially analysed tracer transit after 45 minutes and observed a trend for delay in infected mice during the acute but not the chronic phase (Figure 3c). At 3 weeks p.i. there was also significantly increased organ weight (Figure 3d), so we extended analysis at this time point using an increased parasite inoculum and extended the tracer interval time to 60 minutes. Here we observed evidence of significant small intestine transit delay (Figure 3e).

We next assessed colonic transit using a 90 minute interval after the fluorescent tracer feed. Fluorescence transit appeared similar in infected and control mice at 3 and 6 weeks p.i. (Figure 4a). Unlike the timings used to study transit delay in the upper intestinal tract (Figure 3), the method was less reliable to study the colon in isolation because substantial amounts of dye were still present in the small intestine and we could not quantify any dye that was excreted. Nevertheless, large intestine weights were significantly increased in infected mice at 6 and 30 weeks p.i. (Figure 4b) suggesting a site-specific dysfunction. We therefore employed an alternative assay in which mice were fasted for 4 hr prior to analysis of colon lumen contents. *T. cruzi* infected colons showed significantly greater retention of faeces than controls as shown by pellet counts and both wet and dry total faecal weights, ruling out altered water absorption as an explanation (Figure 4c). The colon-localised transit delay phenotype was highly significant at 6 weeks p.i. and endured into the chronic phase, at 30 weeks p.i. (Figure 4c). By varying the fasting time (0, 2 and 4 hr) we showed that this phenotype was maintained irrespective of stomach fullness and showed distal colon faecal impaction developing in *T. cruzi* infected mice within this timeframe (Figure 4d). The other *T. cruzi* strains exhibiting signs of total GI transit delay in the carmine assay (SN3, Peru, CLBR) also showed significant retention of faeces after 4 hr fasting, whereas strains with normal carmine transit times did not (Supplementary Figure 4). Thus, when GI transit dysfunction occurs in murine chronic *T. cruzi* infections it is predominantly localised to the colon.

**Figure 4:**
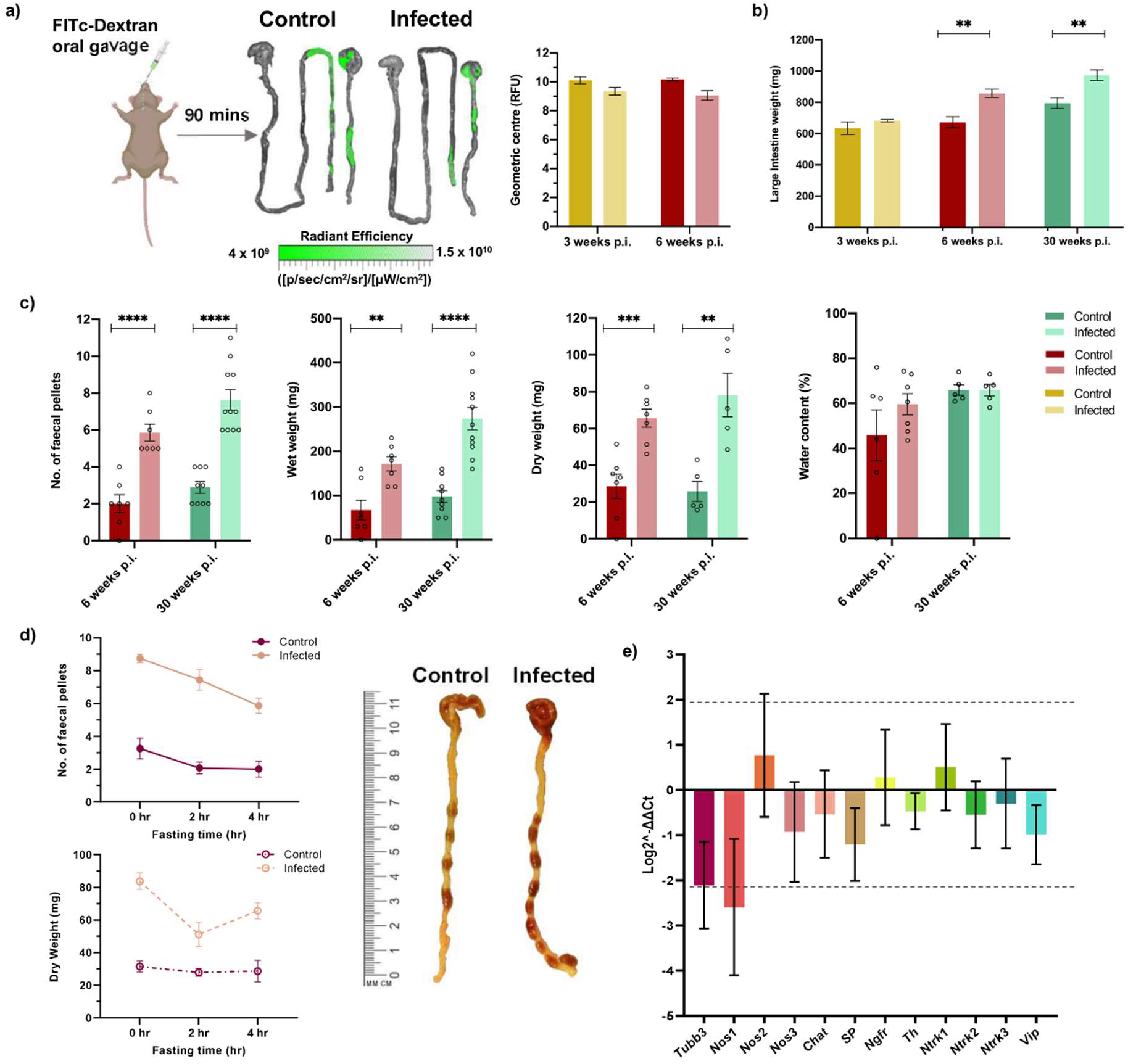
Evidence of colonic transit dysfunction in experimental digestive Chagas disease model. **a)** Schematic diagram of a mouse receiving oral gavage of a green fluorescent marker, 70-kDa FITc-dextran, 90 min prior to termination to trace large intestine transit delay during infection. Representative images of stomach, small and large intestine are superimposed with traces of 70-kDa FITc-dextran travelling through small into large intestine to show transit difference between control and TcI-JR C3H/HeN infected mice. Linear-scale pseudocolor heat map shows minimum and maximum fluorescence intensity of 70-kDa FITc-dextran. Bar plots show quantification of 70-kDa FITc-dextran fluorescence in the large intestine of mice at 3 (*n* = 4 per group) and 6 (*n* = 4 per group) weeks post-infection (p.i.). Fluorescence is expressed as geometric centre which is centre mass of the marker. **b)** Bar plot shows post-mortem weights of large intestine at 3 (*n* = 4 per group), 6 (*n* = 7 per group) and 30 (control *n* = 9, TcI-JR *n* = 11) weeks p.i. **c)** Faecal output analyses between control and TcI-JR C3H/HeN infected mice are expressed as faecal pellet count, wet and dry weight, and percentage of water content at 6 (*n* = 7 per group) and 30 weeks p.i. (*n* = 5-11 per group). **d)** Quantification of the effect of different fasting times on faecal output of mice: number of faecal pellets (*n* = 4-16 per group) and dry faecal weight (*n* = 4-7 per group). Images of mouse large intestine showing faecal impaction during infection at 30 weeks p.i. after 4 hours fasting compared to control. Scale bar is in cm and mm. Data are expressed as mean ± SEM. Statistical significance was tested using unpaired two-tailed Student’s t test (\*\**P* < 0.01; \*\*\**P* < 0.001, **** *P* < 0.0001). **e)** RT-qPCR analysis show log2-fold change in RNA expression of neuronal specific markers: *Tubb3, Nos1, Nos2, Nos3, Chat, SP, Ngfr, Th, Ntrk1, Ntrk2, Ntrk3* and *Vip* in the colon tissue of C3H/HeN naïve control and TcI-JR infected mice (*n* = 5 per group, biological replicates). Data are expressed as Log2^-ΔΔCt^ ± SD. Dashed line represents mean ± 2SD based on distribution of naïve group values.

To further investigate whether the observed functional constipation phenotype was accompanied by alterations at the molecular level, we used RT-qPCR to measure transcript abundance for 12 neuronal and inflammatory response genes in colon tissue from chronically infected mice (Figure 4e). Neuron-specific tubulin β-3 (*Tubb3*) and neuronal nitric oxide synthase (*Nos1*) genes were strongly downregulated by ~75% compared to naïve control mice. Expression of excitatory substance P and inhibitory vasoactive intestinal peptide (*Vip*) ENS neurotransmitters was also decreased, but to a lesser extent. No evidence of altered transcript abundance was found for markers of other enteric neuronal subtypes, tyrosine hydroxylase (*Th*) and acetylcholine (*Chat*), tropomyosin receptor kinases (*Ntrk1*/2/3) or nerve growth factor (*Ngfr*). Taken together, these data indicate a possible downregulation of the enteric nitrergic transmission associated with GI dysfunction in DCD mice, recapitulating observations in human Chagas megasyndromes as well as other enteric neuropathies ^19–23^.

### Chronic infection foci and enteric neuronal damage at organ and tissue micro-domain scales

Our next aim was to investigate disease pathogenesis in this model and commonalities with human DCD. Colon tissue from TcI-JR chronically infected mice (> 210 days p.i.) contained significant lymphocytic inflammatory infiltrates that were diffusely and focally distributed in the smooth muscle layers (Figure 5a). Immunohistochemical labelling of the nerve plexuses within the muscle layers showed that the total amount of neuron-specific tubulin (TuJ1) protein within myenteric ganglia was lower on average in infected mice, but this was not statistically significant (Figure 5b). However, there was a conspicuous spatial disorganisation of TuJ1 in a subset of ganglia, associated with the appearance of anomalous internal acellular structures in these ganglia, which were refractory to common histological dyes (Figure 5b, Supplementary Figure 5). To investigate this with greater precision, we used whole mount immunofluorescence analysis of the neuronal cell body marker HuC/D. This revealed a dramatic loss of neurons across the proximal, mid and distal colon myenteric plexus (Figure 5c, 5d). This was not a product of a reduced number of ganglia (Figure 5e), rather a highly significant reduction in neurons per ganglion (Figure 5f).

**Figure 5:**
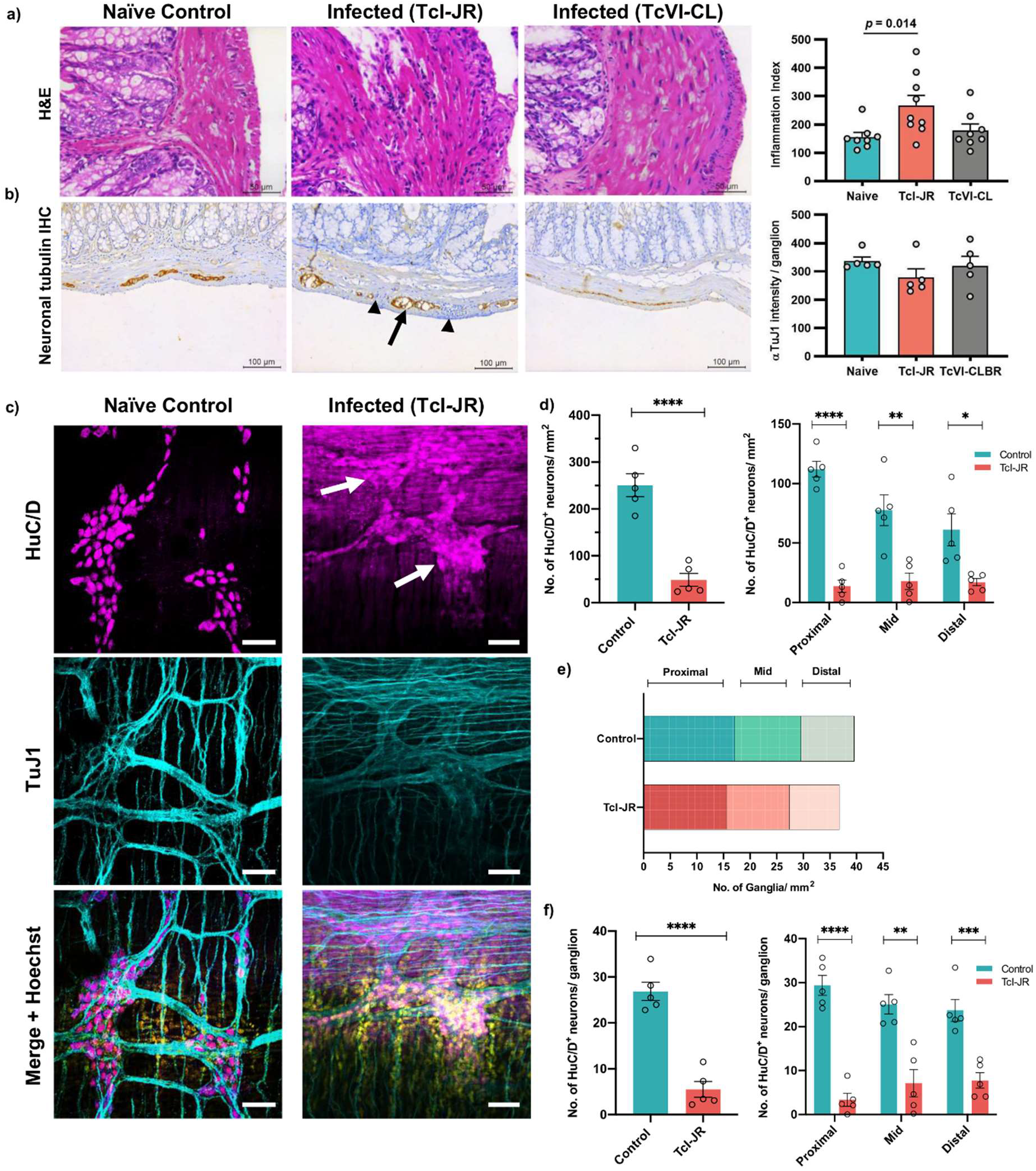
Effects of chronic *T. cruzi* infection on the enteric nervous system. **a)** Representative brightfield images of 5 μm thick colon sections stained with haematoxylin-eosin. Images were taken at 100x magnification, scale bar = 50 μm. Adjacent bar plot shows number of nuclei per field to quantify cellular infiltration in TcI-JR (*n* = 8), TcVI-CLBR (*n* = 10) infected mice compared to naïve controls (*n* = 8). **b)** Representative brightfield images of 5 μm thick colon sections to detect change in pathology during *T. cruzi* infection detected by immunohistochemistry. Images were taken at 100x magnification, scale bar = 50 μm. Adjacent bar plot shows percentage of neuronal tubulin (Tuj1) immunoreactivity in naïve control (*n* = 8), TcI-JR-(*n* = 8) and TcVI-CLBR (*n* = 9) infected mice. **c)** Representative immunofluorescent confocal images of whole-mount colon samples to show the change in anti-HuC/D stained neuronal cell bodies (magenta, top panel) and anti-Tuj1 stained neural network (cyan, middle panel) in the myenteric plexus during *T. cruzi* infection. Bottom panel shows merged images with DAPI nuclei stain (yellow). White arrows indicate damaged ganglionic neuronal cell bodies. Images were taken at 40x magnification, scale bar: 50 μm. **d)** Bar plots show number of HuC/D positive neuronal cell bodies per field of view in naïve control and TcI-JR infected whole colon samples (left) and from selected regions of the colon: proximal, mid and distal (right; *n* = 5 per group, all). **e)** Quantification of number of ganglia in naïve control and TcI-JR infected samples from proximal, mid and distal colon (*n* = 5 per group). **f)** Bar plots show number of HuC/D positive neuronal cell bodies per ganglion in naïve control and TcI-JR infected whole colon samples (left) and from selected regions of the colon: proximal, mid and distal (right; *n* = 5 per group, all). All data and images are obtained from matched naïve control and infected mice at 30 weeks post-infection. Data are expressed as mean ± SEM. Statistical significance was tested using unpaired two-tailed Student’s t test (\**P* < 0.05; \*\**P* < 0.01; \*\*\**P* < 0.001, **** *P* < 0.0001).

A critical question for rational design of therapies for DCD is whether *T. cruzi* and the associated host response continues to drive ENS pathology during the chronic phase of infection. At this stage, very few colon cells are infected at any one time and parasite foci are spatiotemporally dynamic, with an intracellular lytic cycle lasting 1-2 weeks before motile trypomastigotes migrate within and between tissues ^24^. Thus, any temporal association between infection and ENS damage is likely highly localised and rare at any snapshot in time. Indeed, there was no correlation between chronic endpoint parasite loads in colon regions and the level of local denervation (Figure 6a). We also observed both denervated and intact myenteric ganglia immediately adjacent to each other (Figure 6b). Using dual bioluminescent-fluorescent reporter parasites ^25^ we were able to visualise rare chronic infection foci at single cell resolution. In most cases, infected cells were early in the proliferative cycle, with 10-50 amastigote forms, and they were located in close proximity to intact enteric nerve fibres (Figure 6c, Supplementary Figure 6). We also captured a very rare, mature pseudocyst at the point of rupture, with thousands of intracellular parasites and trypomastigote forms escaping the site (Figure 6d). The ENS at the level of this pseudocyst was almost completely ablated, whereas the overlying and laterally adjacent ENS networks were intact (Figure 6e). Taken together, our data demonstrate there is an enduring association, at a highly localised tissue micro-domain scale, between chronic parasitism of the gut wall and ENS lesions.

**Figure 6:**
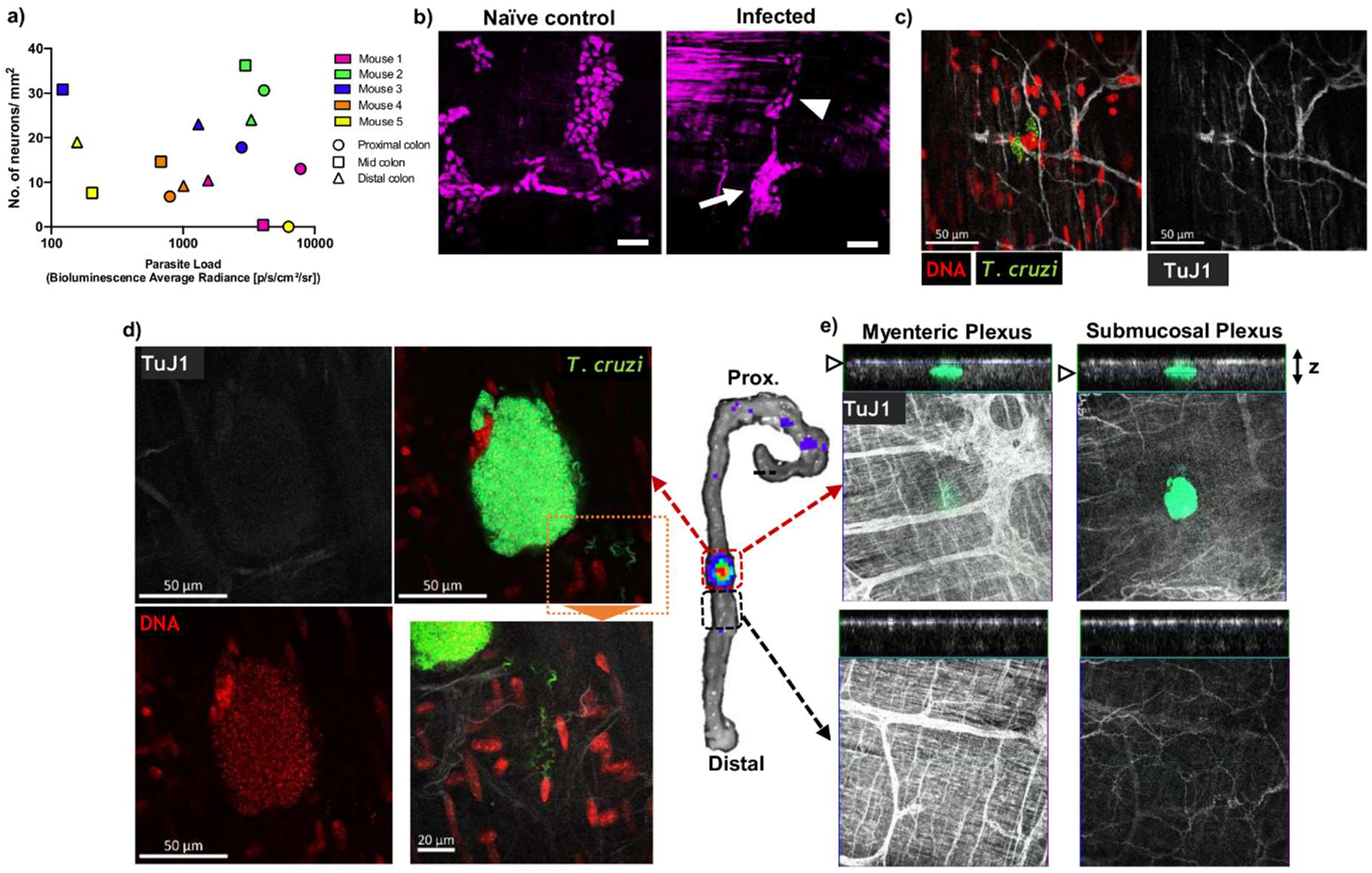
Chronic phase colonic *T. cruzi* infection foci and ENS ablation at the tissue micro-domain scale. **a)** Lack of correlation between end-point colon parasite loads measured by *ex vivo* bioluminescence intensity and degree of colon myenteric plexus denervation. **b-e)** Whole mount (immuno)fluorescence analysis of colonic muscularis from C3H mice chronically infected with *T. cruzi.* **b)** HuC/D+ neuronal cell bodies in colonic myenteric ganglia. Naïve control shows normal morphology; infected mice exhibit adjacent ganglia with both intact (arrowhead) and disrupted (arrow) staining patterns. **c)** Imaging individual *T. cruzi (*mNeonGreen^*+*^) infected cells at early stage of parasite replication cycle adjacent to intact enteric neuron fibres (TuJ1^+^). **d-e)** Bioluminescence *ex vivo* image (centre)-guided analysis of parasitized and parasite-free tissue micro-domains. **d)** Mature parasite pseudocyst containing >1000 flagellated trypomastigotes with extracellular trypomastigotes in the local tissue parenchyma (inset) with faint neuronal (TuJ1) staining. **e)** Z-stack slices at the level of myenteric and submucosal neuronal plexuses showing highly localised loss of TuJ1 staining around the rupturing parasite pseudocyst.

## Discussion

Understanding of the mechanism of DCD pathogenesis remains rudimentary and a lack of experimental tools hampers progress. Post-mortem and biopsy studies of human DCD cases found reduced numbers of enteric neurons and glial cells ^26–28^. These are important insights into late and terminal disease states, but they provide limited information on pathogenesis and relationships with infection load or distribution. *T. cruzi* infected mice do not develop digestive megasyndromes, but these are late stage manifestations of human disease, and usually take many years to develop. Nevertheless, denervation and other features of nascent enteropathy have been described in mouse models at the histological level ^14,29–32^. Delayed transit has also been reported^33,34^ but neither the GI region involved nor associations with infection dynamics were determined. In this study, we present new experimental chronic *T. cruzi* infection models that, crucially, feature co-localised parasite persistence, enteric denervation and delayed transit as a key functional symptom of DCD. This will now enable pre-clinical drug development to focus on this form of CD, supported by an ability to undertake longitudinal monitoring of individual animal parasite loads using bioluminescence imaging.

In keeping with human disease epidemiology ^35^, our data show that both host and parasite genetics contribute to murine DCD susceptibility. The digestive tract in mice is a universal reservoir of chronic infection, but only a few *T. cruzi* strains caused transit dysfunction within the timeframe of these experiments. Disease severity was also higher in C3H/HeN than BALB/c mice, a finding previously observed for murine cardiac CD ^9^, and is consistent with the heterogeneous clinical outcomes observed in humans ^36^. Thus, gut parasitism alone is not sufficient as an explanation for DCD development. Investigation of parasite virulence and variability in the host immune response will be required to gain further insight into the determinants of susceptibility and resistance.

Our results challenge the prevailing theory that DCD is a result of collateral damage to the ENS, resulting specifically from the acute inflammatory response against *T. cruzi* ^5,37^. This idea was rooted in an inability to detect gut-localised parasites in chronic infections, which has only recently been overcome by the development of highly sensitive bioluminescence imaging methods ^8,9^. By combining live parasite imaging and gut tracer analyses, we found enduring associations between infection of the colon and local transit impairment at >6 months post-infection, and moreover at the tissue micro-domain scale between single infected cells and ENS lesions. Treatment of human chronic infections with anti-parasitic chemotherapy (benznidazole or nifurtimox) may therefore be beneficial to prevent or alleviate DCD. Outstanding questions include whether the patchiness of ENS damage is explained by the stochastic distribution of parasites, or because particular subsets of ganglia or neurons differ in susceptibility, and if so, why. We focussed on analysis of neurons in the myenteric plexus, but it will be important to explore other ENS components, including potential regulatory or neuroprotective functions of enteric glial cells ^38^ and broader factors known to influence neuro-immune interactions in the gut, such as host metabolism ^39^ and microbiota ^40^.

## Materials and Methods

### Parasites and infections

Transgenic clones of *T. cruzi* TcI-JR and TcVI-CLBR constitutively expressing the red-shifted firefly luciferase variant *PPy*RE9h ^15^ alone or fused to mNeonGreen were described previously ^9,25^. Equivalent clones for other *T. cruzi* strains were generated by transfection of the DNA construct pTRIX2-RE9h (TcI-C8, TcI-X10/4, TcIII-Arma18, TcVI-Peru), or by cas9-mediated replacement of the LucRE9h gene with dual reporters, namely LucRE9h::Neon (TcVI-CLBR, TcII-Pot7a, TcIV-X10610) and LucRE9h::mScarlet (TcI-ArePe, TcI-SN3), using the T7 RNA polymerase/cas9 system ^25^. *In vitro* epimastigotes were cultivated in supplemented RPMI-1640 medium at 28°C with 150 μg ml^−1^ G418 or hygromycin B, 5 μg ml^−1^ puromycin or 10 μg ml^−1^ blasticidin as appropriate. Metacyclic trypomastigotes (MTs) from stationary phase cultures were used to infect MA104 monkey kidney epithelial cell monolayers in MEM media + 5% FBS at 37°C and 5% CO_2_. Tissue culture trypomastigotes (TCTs) were obtained from the supernatant of infected cells after 5 to 21 days, depending on the parasite strain.

### Animals and infections

All *in vivo* experiments were performed in accordance with UK Home Office regulations under the Animal Scientific Procedure Act (ASPA) 1986, project license 70/8207 or P9AEE04E, and were approved by LSHTM Animal Welfare Ethical Review Board. Female BALB/c and C3H/HeN mice, postnatal days 42-56, were purchased from Charles River (UK). Female CB17 SCID mice were bred in-house. All mice were housed on a 12 hr light/dark cycle, with food and water available *ad libitum* unless otherwise stated. Mice were maintained under specific pathogen-free conditions in individually ventilated cages.

SCID mice were infected with up to 5 × 10^5^ *in vitro*-derived TCTs in 0.2 ml PBS via i.p. injection. All infected SCID mice developed fulminant infections and were euthanised at or before humane end-points. Blood trypomastigotes (BTs) were derived from parasitaemic SCID mouse blood directly or after enrichment, achieved by allowing blood samples to settle for 1 hr at 37° C. BALB/c and C3H mice were infected by i.p injection of 10^3^ or 10^4^ BTs or TCTs depending on the experiment.

At experimental end-points, mice were sacrificed by ex-sanguination under terminal anaesthesia (Euthatal/Dolethal 60 mg kg^−1^, i.p.) or by cervical dislocation. Organs and tissues of interest were excised, imaged (see below) and either snap-frozen, fixed in 10% Glyofixx or transferred to ice-cold DMEM media. The weight of organs and tissues of interest were recorded.

### Total GI transit time assay

Mice were gavaged p.o. with 200 μl of 6% w/v Carmine red dye solution in 0.5% methyl cellulose mixed in distilled water and returned to their home cage. After 75 min, mice were individually separated into containers and the time of excretion of the first red faecal pellet was recorded. A maximum assay cut-off time of 4 hr was implemented. Total intestinal transit time was calculated as the time taken from gavage to output of the first red pellet.

### *In vivo* bioluminescence imaging

Mice were injected with 150 mg kg^−1^ d-luciferin i.p., then anaesthetised using 2.5% (v/v) gaseous isoflurane in oxygen. After 10-20 min, bioluminescence imaging was performed using an IVIS Lumina II or Spectrum system (PerkinElmer), with acquisition time and binning adjusted according to signal intensity. Mice were revived and returned to cages after imaging. To estimate parasite burden in live mice, regions of interest (ROIs) were drawn to quantify bioluminescence expressed as total flux (photons/second) ^8,9^. The detection threshold was determined using uninfected control mice. All bioluminescence data were analysed using LivingImage v4.7.3.

### *Ex vivo* bioluminescence imaging

Mice were injected with 150 mg kg^−1^ d-luciferin i.p. 5-7 min before euthanasia. Trans-cardiac perfusion was performed with 10 ml of 0.3 mg ml^−1^ d-luciferin in PBS. Tissues and organs of interest (typically lymph nodes, heart, spleen, skeletal muscle, GI tract and associated mesenteries) were collected and soaked in PBS containing 0.3 mg ml^−1^ d-luciferin. Bioluminescence imaging was performed as above. To quantify parasite load as a measure of infection intensity, bioluminescence was calculated by outlining region of interest (ROI) on each sample and expressed as radiance (photons second^−1^ cm^−2^ sr^−1^). Fold change in radiance was determined by comparing samples from infected mice with the equivalent tissues from uninfected, age-matched control mice. To determine the detection threshold, fold change in radiance of an empty ROI on images from infected mice were compared with matching empty ROI on images from uninfected controls ^8^.

### Fluorescent tracer transit assay

Mice were fasted (or not) for 2 or 4 hr before euthanasia. They were administered 70-kDa FITC dextran (100 μl, 5 mg ml^−1^ d.H_2_O) or Rhodamine dextran (100 μl, 10mg ml^−1^ d.H_2_O) by oral gavage 5, 45 or 90 min before euthanasia to target the stomach, small or large intestine transit, respectively. As an extension of the *ex vivo* bioluminescence necropsy (see above), fluorescence images were obtained using excitation filters set at 465/535 nm and emission filters at 502/583 nm for FITC/Rhodamine (f-stop: 16, exposure: 2 s). The relative fluorescence of the tracers was measured from the images by drawing ROIs using LivingImage 4.7.3 software. The GI tract images starting from the stomach to the colon were cut digitally in 15 equal segments and the centre of mass (geometric centre) of the signals were determined. The geometric centre was calculated using the following equation, GC = ∑ (% of total fluorescent signal per segment * segment number) / 100) ^41^.

### Faecal analyses

The colon tissue was separated and cleaned externally with PBS. The faecal pellets were gently removed from the lumen of the colon, counted and the combined wet weight was recorded. The faecal pellets were collected into a 12-well plate and left to dry in a laminar flow cabinet overnight. The dry weight was then recorded and the percent water content was estimated as the difference between wet and dry weights.

### Histopathology and Immunohistochemistry

Paraffin-embedded, fixed tissue blocks were prepared and 3-5 μm sections were stained with haematoxylin and eosin as described ^9,42^. For tubulin β-3 immunohistochemistry, sections were subjected to heat-induced epitope retrieval by incubation in 10 mM sodium citrate, 0.05% Tween20 for 30 min then cooled and rinsed in distilled water. Sections were blocked with 10% sheep serum and 1% BSA in TBS for 30 min then incubated at 4°C overnight with 1 μg ml^−1^ rabbit polyclonal anti-tubulin β-3 IgG (Biolegend) and 1% BSA in TBS. Sections were then washed with 0.025% Triton X-100 in TBS and endogenous peroxidase activity was quenched with 3% H_2_O_2_ for 30min. Bound primary antibody was labelled with excess volume of HRP polymer anti-rabbit IgG reagent (Vector Labs) with 1% BSA in TBS for 30 min. Slides were then washed as previously and incubated with DAB (Thermo) for 5 min. Sections were counterstained with haematoxylin and mounted with DPX.

Images were acquired using a Leica DFC295 camera attached to a Leica DM3000 microscope. For analysis of inflammation, nuclei were counted automatically using the Leica Application Suite V4.5 software (Leica). DAB intensity was analysed as integrated density in ImageJ.

### Immunofluorescence analysis

Colon tissues were transferred into ice-cold DMEM after necropsy. Tissues were cut open along the mesentery line, rinsed with PBS, then stretched and pinned on Sylgard 184 plates. The mucosal layer was peeled away using forceps under a dissection microscope and the remaining muscularis wall tissue was fixed in paraformaldehyde (4% w/v in PBS) for 45 min at room temperature. Tissues were washed with PBS for up to 45 min at room temperature and permeabilised with PBS containing 1% Triton X-100 for 2 hr, followed by blocking for 1 hr (10% sheep serum in PBS containing 1% Triton X-100). Tissues were incubated with primary antibodies (mouse anti-HuC/D IgG clone 16A11 at 1:200 [Thermofisher], rabbit polyclonal anti-tubulin β-3 IgG at 1:500 [Biolegend]) in PBS containing 1% Triton X-100 for 48 h at 4 °C. Tissues were washed with PBS, then incubated with secondary IgG (goat anti-mouse Alexa546, goat anti-rabbit Alexa633, both 1:500, ThermoFisher) in PBS containing 1% Triton X-100 for 2 h and counterstained with Hoechst 33342 (1:10 000) at room temperature. To assess antibody specificity, control tissues were incubated without the primary antibody. Tissues were mounted on glass slides using FluorSave mounting medium (Merck).

Whole mounts were examined and imaged with a LSM880 confocal microscope using a 40x objective (Zeiss, Germany). Images were captured as Z-stack scans of 21 digital slices with interval of 1 μm optical thickness. Five Z-stacks were acquired per region (proximal, mid and distal colon), per animal. Cell counts were performed on Z-stacks after compression into a composite image using the cell counter plug-in of FIJI software. Neuronal density was calculated as the number of HuC/D^+^ neuron cell bodies per field of view. HuC/D signal was associated with high background outside ganglia in samples from infected mice, attributed to binding of the secondary anti-mouse IgG to endogenous IgG, so ENS-specific analysis was aided by anti-TuJ1 co-labelling and assessment of soma morphology. The number of intact ganglia in each myenteric plexus image was also counted, along with number of HuC/D^+^ neurons per ganglia.

### RT-qPCR

Colon tissue samples were snap frozen on dry ice and stored at −70°C. For RNA extraction, samples were thawed and homogenised in 1 ml Trizol (Invitrogen) per 30-50 mg tissue using a Precellys 24 homogeniser (Bertin). To each sample, 200 μl of chloroform was added and mixed by vortex after which the phases were separated by centrifugation at 13,000 g at 4°C. RNA was extracted from the aqueous phase using the RNeasy Mini Kit (Qiagen) with on-column DNAse digestion as per manufacturer’s protocol. The quantity and quality of RNA was assessed using Qubit Fluorimeter (Thermofisher). cDNA was synthesised from 1 μg of total RNA using Superscript IV VILO mastermix (Invitrogen), as per manufacturer’s protocol, in reaction volumes of 20 μl. A final cDNA volume of 100 μl was made by adding RNase-free DEPC water (1:5 dilution) and stored at −20 °C until further use. qPCR reactions contained 4 μl of cDNA (1:50 dilution) and 200 nM of each primer (supplementary table 1) and QuantiTect SYBR Green PCR master mix (Qiagen) or SensiFAST SYBR Hi-ROX kit (Bioline). Reactions were run using Applied Biosystems 7500 fast RT-PCR machine (Thermofisher) as per manufacturer’s protocol. Data were analysed by the ΔΔCt method ^43^ using murine *Gapdh* as the endogenous control gene.

### Statistics

Individual animals were used as the unit of analysis. No blinding or randomisation protocols were used. Statistical differences between groups were evaluated using unpaired two-tailed Student’s t-test or one-way ANOVA with Tukey’s post-hoc correction for multiple comparisons. Pearson correlation analyses was used to evaluate relationships between variables. These tests were performed in GraphPad Prism v.8 or R v3.6.3. Differences of *p* < 0.05 were considered significant.

---

**Supplementary Figure 1.**
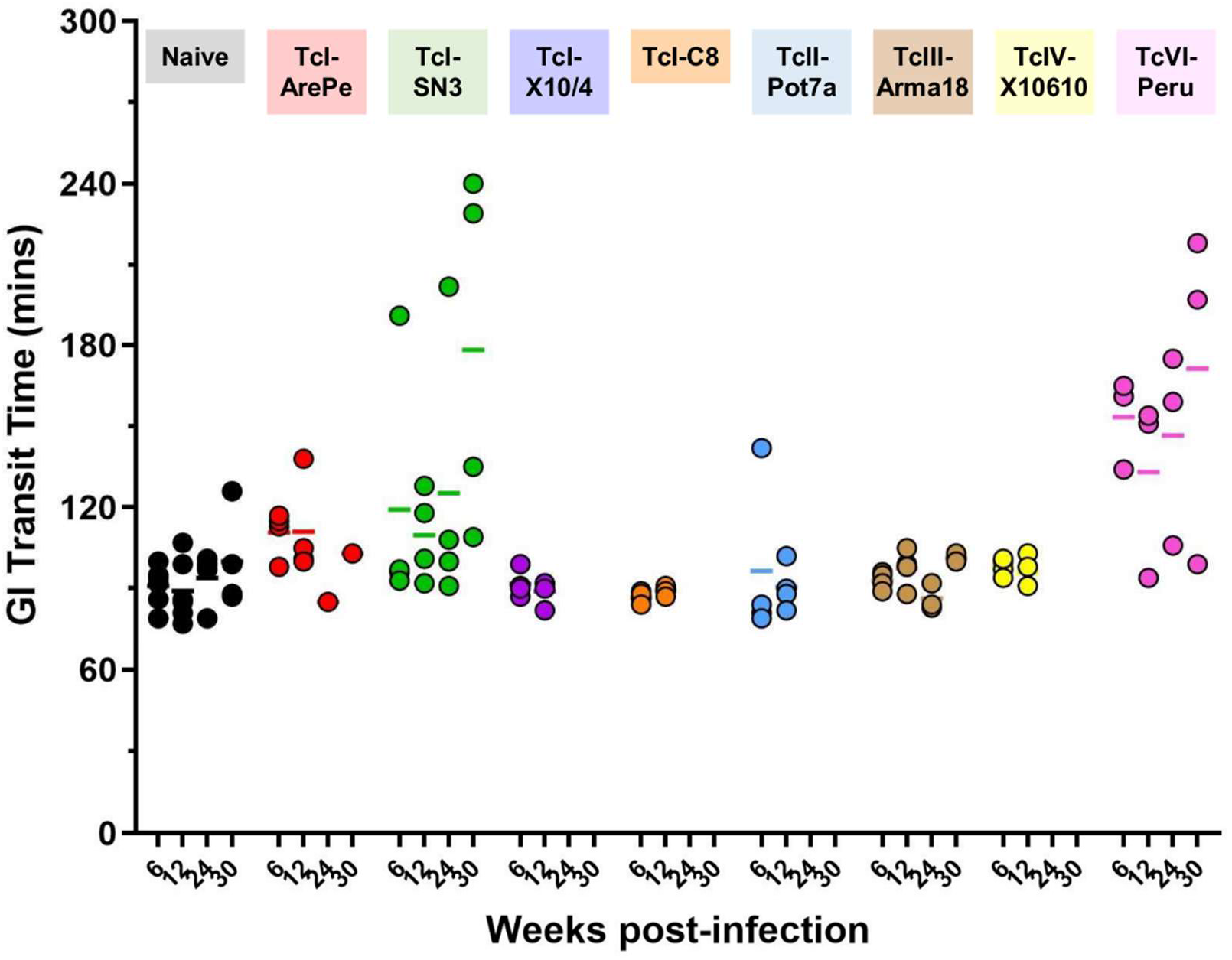
Gastrointestinal transit dysfunction screen in mice infected with different strains of *T. cruzi*. Data are gastrointestinal (GI) transit time at indicated weeks post-infection (p.i.) for C3H/HeN mice in the following infection groups: naive control (n = 4-6), TcI-ArePe (n = 1-4), TcI-SN3 (n = 4), TcI-SylvioX10/4 (n = 4) and TcI-C8 (n = 4), TcII-Pot7a (n = 4), TcIII-Arma18 (n = 3-4), TcIV-X10610 (n = 4) and TcVI-Peru (n = 3).

**Supplementary Figure 2.**
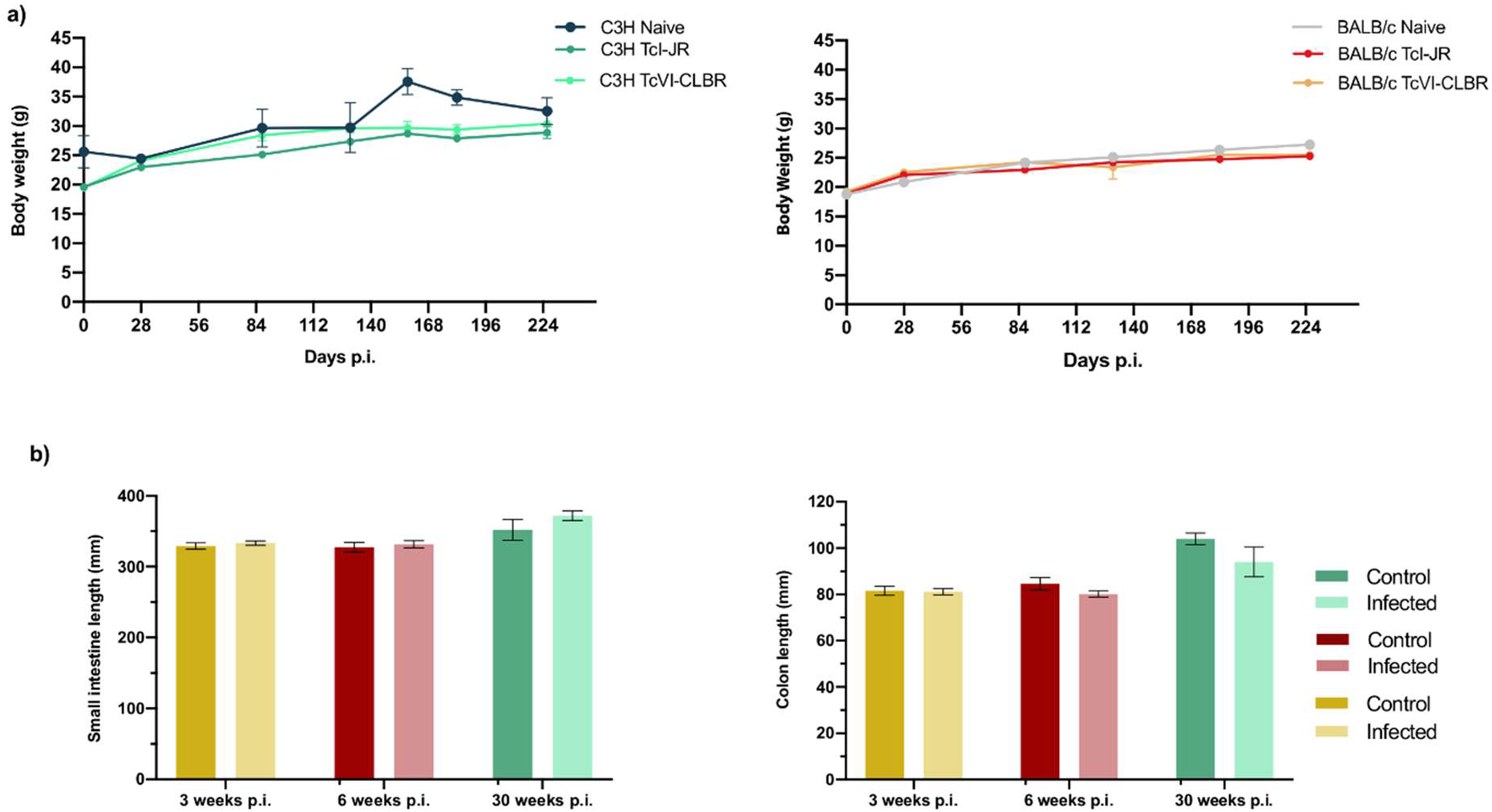
Anatomical measures of gastrointestinal *T. cruzi* infection mouse models. **a)** Body weights of naïve control (*n* = 3-10), TcI-JR (*n* = 5– 22) and TcVI-CLBR (*n* = 5-20) infected C3H/HeN (left) and BALB/c mice (right) vs. days post-infection (p.i.). **b)** Bar plots show length of small intestine and colon of control and TcI-JR C3H/HeN mice at 3 (*n* = 24 per group), 6 (*n* = 27 per group) and 30 (*n* = 5 per group) weeks p.i. Data are expressed as mean ± SEM.

**Supplementary Figure 3.**
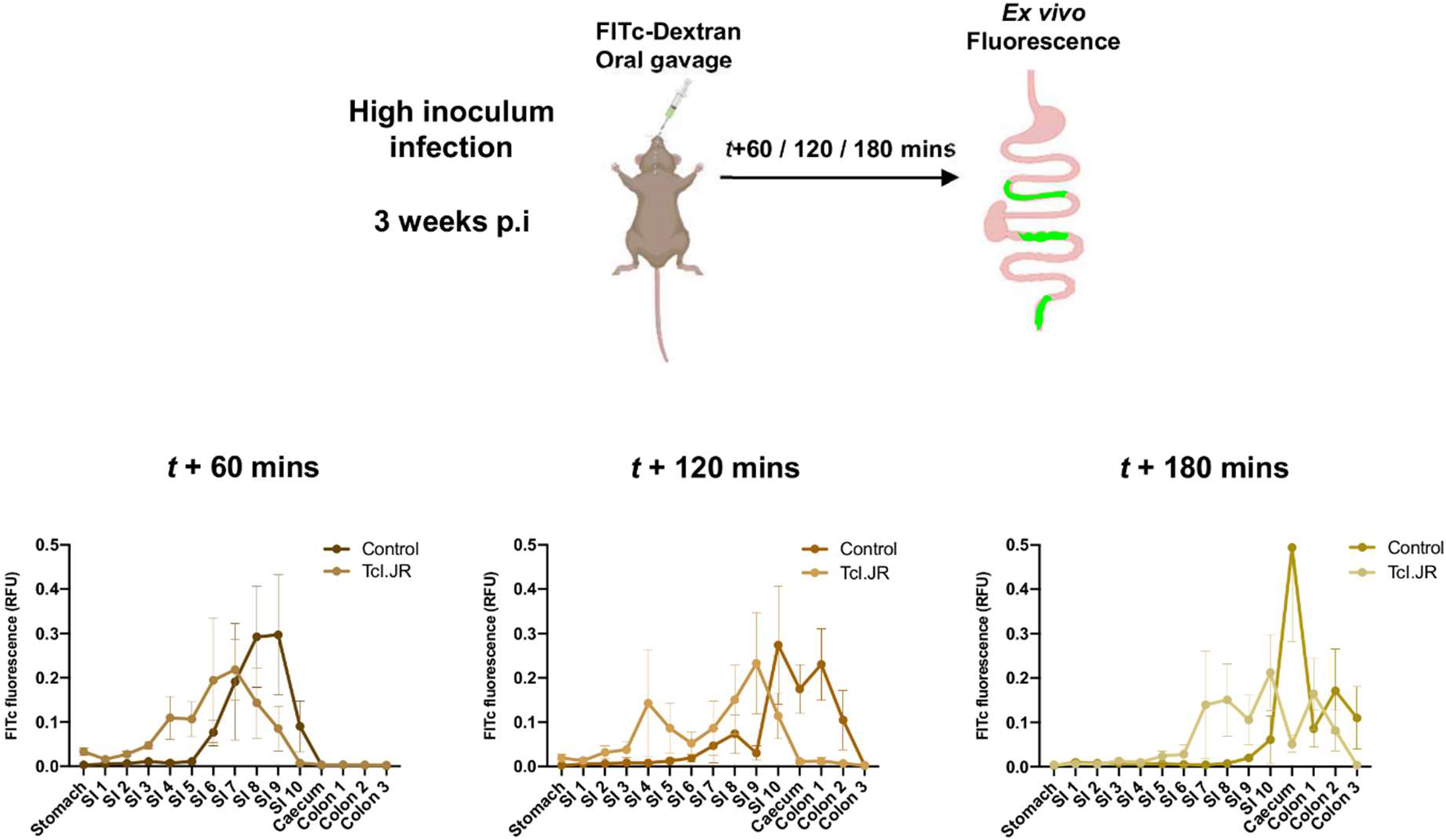
Fluorescent tracer imaging assay for gastrointestinal (GI) transit in high acute *T. cruzi* infection. Schematic diagram of a mouse receiving oral gavage of a green fluorescent marker, 70-kDa FITc-dextran, 60 or 120 or 180 minutes prior to termination to trace localised GI transit delay during acute infection. Quantification of 70-kDa FITc-dextran fluorescence in different parts of the GI tract (S1-S210: small intestine scored into 10 equal sections) of naïve control and TcI-JR C3H/HeN (*n* = 4 per group) mice at 3 weeks post-infection. All mice in this experiment were infected with a high inoculum of TcI-JR parasites. Data are expressed as mean ± SEM.

**Supplementary Figure 4.**
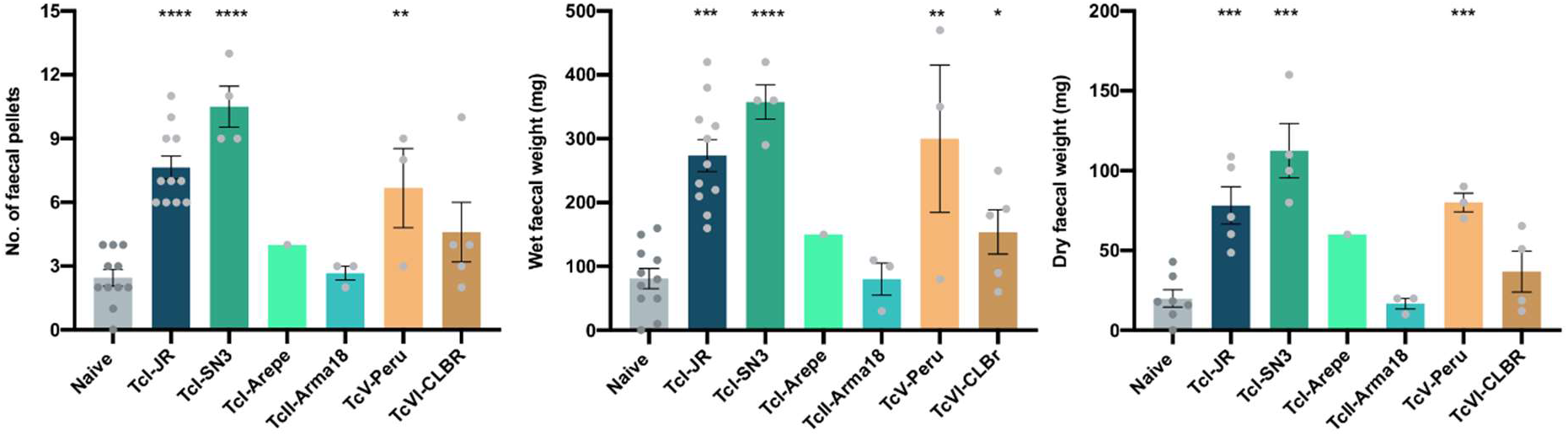
Comparison of colonic transit dysfunction in different models of experimental digestive Chagas disease. Faecal output analyses are expressed as faecal pellet count, wet and dry pellet weight at 30 weeks post-infection (p.i.) in the following groups: naive control (*n* = 7-11), TcI-JR (*n* = 5-11), TcI-SN3 (*n* = 4), TcI-ArePe (*n* = 1), TcIII-Arma18 (*n* = 3), TcVI-Peru (*n* = 3) and TcVI-CLBR (*n* = 4-5).

**Supplementary Figure 5.**
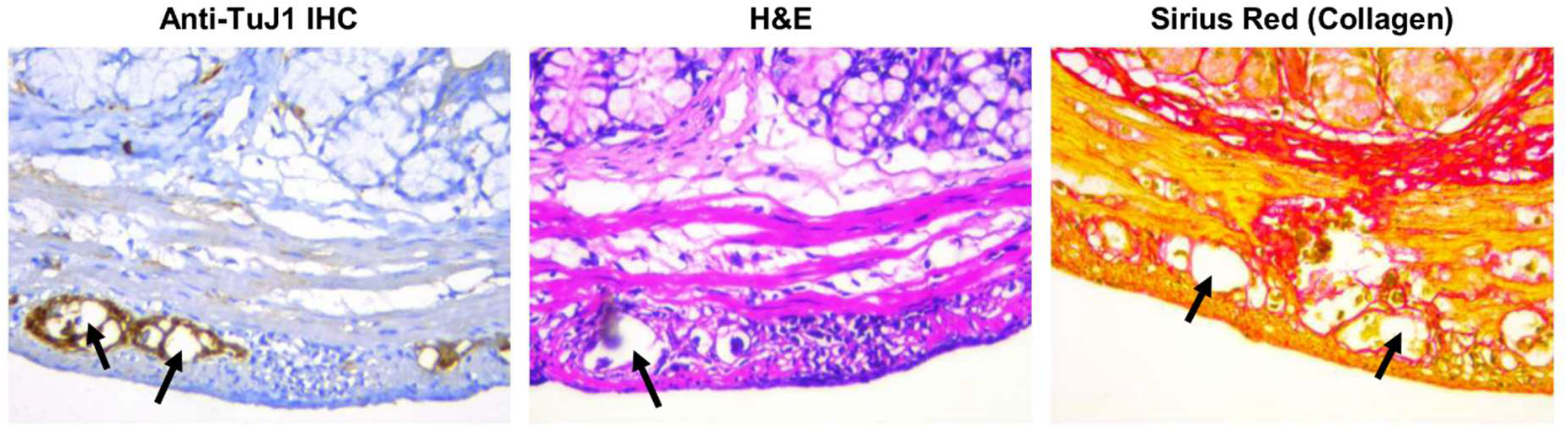
Myenteric neuronal plexus lesions in *T. cruzi* infection. Acellular structures (arrows) within myenteric plexus ganglia from C3H mice with chronic TcI-JR infections that are refractory to staining by the indicated method.

**Supplementary Figure 6.**
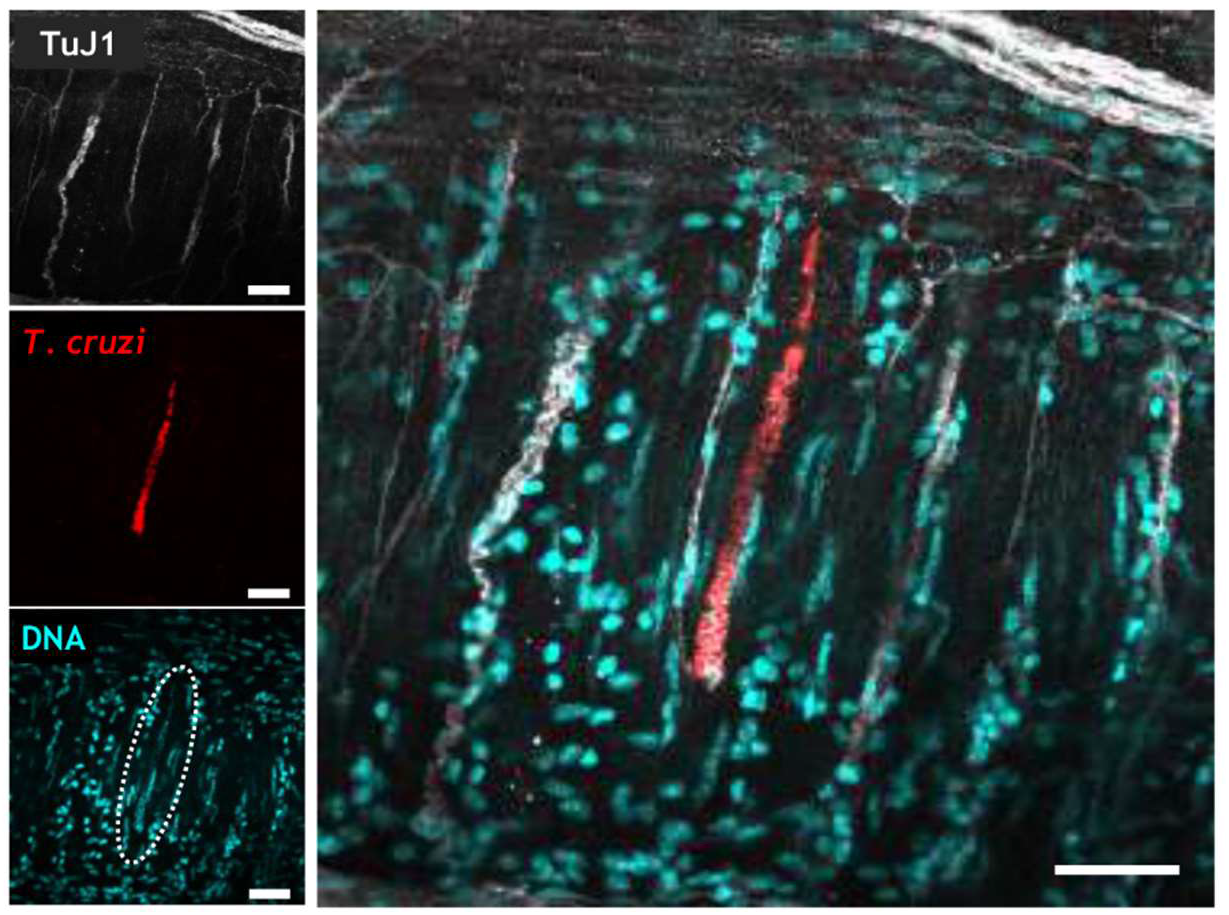
Chronic phase colonic *T. cruzi* infection foci of TcI-SN3 parasites in the ENS. Representative immunofluorescent z-stack confocal images of whole-mount colonic muscularis from C3H mice chronically infected with fluorescent TcI-SN3 (scarlet+) parasites. Image shows the localisation of TcI-SN3 parasites (red) in the submucosal layer of the ENS stained with anti-Tuj1 (phase). DAPI (cyan) shows nuclei stain and white circle indicates DNA of the parasite nest. Images were taken at 40x magnification, scale bar: 50 μm.

## Acknowledgements

We thank Hernán Carrasco, Michael Miles, Manuel Sánchez-Moreno, Omar Triana and Matthew Yeo for sharing parasite strains and the LSHTM Biological Services Facility staff for technical support and animal husbandry. Some figure panels were created with BioRender.com. The work was funded by an MRC New Investigator Research Grant (MR/R021430/1) and an EU Marie Curie Fellowship (grant agreement no. 625810).

## References

1 Rassi Jr, A., Rassi, A. & Marin-Neto, J. A. Chagas disease. Lancet 375, 1388–1402 (2010).

2 Iantorno, G. et al. The enteric nervous system in chagasic and idiopathic megacolon. Am J Surg Pathol 31, 460–468 (2007).

3 Meneghelli, U. G. Chagasic enteropathy. Revista da Sociedade Brasileira de Medicina Tropical 37, 252–260 (2004).

4 Bern, C. Antitrypanosomal Therapy for Chronic Chagas’ Disease. New Engl J Med 364, 2527–2534, doi:doi:10.1056/NEJMct1014204 (2011).

5 Köberle, F. Chagas’ disease and Chagas’ syndromes: the pathology of American trypanosomiasis. Adv Parasitol 6, 63–116 (1968).

6 de Oliveira, R. B., Troncon, L. E., Dantas, R. O. & Menghelli, U. G. Gastrointestinal manifestations of Chagas’ disease. Am J Gastroenterol 93, 884–889 (1998).

7 Arantes, R. M. et al. Interferon-gamma-induced nitric oxide causes intrinsic intestinal denervation in Trypanosoma cruzi-infected mice. Am J Pathol 164, doi:10.1016/s0002-9440(10)63222-1 (2004).

8 Lewis, M. D. et al. Bioluminescence imaging of chronic *Trypanosoma cruzi* infections reveals tissue-specific parasite dynamics and heart disease in the absence of locally persistent infection. Cell Microbiol 16, 1285–1300, doi:10.1111/cmi.12297 (2014).

9 Lewis, M. D., Francisco, A. F., Taylor, M. C., Jayawardhana, S. & Kelly, J. M. Host and parasite genetics shape a link between *Trypanosoma cruzi* infection dynamics and chronic cardiomyopathy. Cell Microbiol 18, 1429–1443, doi:10.1111/cmi.12584 (2016).

10 Laranjeira, C. et al. Glial cells in the mouse enteric nervous system can undergo neurogenesis in response to injury. J Clin Invest 121, 3412–3424, doi:10.1172/JCI58200 (2011).

11 Kulkarni, S. et al. Adult enteric nervous system in health is maintained by a dynamic balance between neuronal apoptosis and neurogenesis. PNAS 114, E3709–E3718, doi:10.1073/pnas.1619406114 (2017).

12 Muller, Paul A. et al. Crosstalk between Muscularis Macrophages and Enteric Neurons Regulates Gastrointestinal Motility. Cell 158, 300–313, doi:https://doi.org/10.1016/j.cell.2014.04.050 (2014).

13 Gabanyi, I. et al. Neuro-immune Interactions Drive Tissue Programming in Intestinal Macrophages. Cell 164, 378–391, doi:https://doi.org/10.1016/j.cell.2015.12.023 (2016).

14 do Carmo Neto, J. R. et al. Correlation between intestinal BMP2, IFNgamma, and neural death in experimental infection with Trypanosoma cruzi. PLoS One 16, e0246692, doi:10.1371/journal.pone.0246692 (2021).

15 Branchini, B. R. et al. Red-emitting luciferases for bioluminescence reporter and imaging applications. Anal Biochem 396, 290–297, doi:10.1016/j.ab.2009.09.009 (2010).

16 Machado, C. A. & Ayala, F. J. Nucleotide sequences provide evidence of genetic exchange among distantly related lineages of Trypanosoma cruzi. Proc Natl Acad Sci U S A 98, 7396–7401, doi:10.1073/pnas.121187198 (2001).

17 Westenberger, S. J., Barnabé, C., Campbell, D. A. & Sturm, N. R. Two hybridization events define the population structure of *Trypanosoma cruzi*. Genetics 171, 527–543, doi:10.1534/genetics.104.038745 (2005).

18 Lewis, M. D. et al. Recent, independent and anthropogenic origins of Trypanosoma cruzi hybrids. PLoS Negl Trop Dis 5, e1363, doi:10.1371/journal.pntd.0001363 (2011).

19 da Silveira, A. B. et al. Neurochemical coding of the enteric nervous system in chagasic patients with megacolon. Dig Dis Sci 52, 2877–2883, doi:10.1007/s10620-006-9680-5 (2007).

20 Nascimento, R. D., Martins, P.R.,, Duarte, G.D. & Reis, D.D. Decrease of Nitrergic Innervation in the Esophagus of Patients with Chagas Disease: Correlation with Loss of Interstitial Cells of Cajal. International Journal of Pathology and Clinical Research 3, 59, doi:10.23937/2469-5807/1510059 (2017).

21 Dickson, E. J., Heredia, D. J., McCann, C. J., Hennig, G. W. & Smith, T. K. The mechanisms underlying the generation of the colonic migrating motor complex in both wild-type and nNOS knockout mice. Am J Physiol Gastrointest Liver Physiol 298, G222–232, doi:10.1152/ajpgi.00399.2009 (2010).

22 McCann, C. J. et al. Transplantation of enteric nervous system stem cells rescues nitric oxide synthase deficient mouse colon. Nat Commun 8, 15937, doi:10.1038/ncomms15937 (2017).

23 Rivera, L. R., Poole, D. P., Thacker, M. & Furness, J. B. The involvement of nitric oxide synthase neurons in enteric neuropathies. Neurogastroenterology & Motility 23, 980–988, doi:https://doi.org/10.1111/j.1365-2982.2011.01780.x (2011).

24 Ward, A. I. et al. *In Vivo* Analysis of *Trypanosoma cruzi* Persistence Foci at Single-Cell Resolution. mBio 11, e01242–01220, doi:10.1128/mBio.01242-20 (2020).

25 Costa, F. C. et al. Expanding the toolbox for *Trypanosoma cruzi*: A parasite line incorporating a bioluminescence-fluorescence dual reporter and streamlined CRISPR/Cas9 functionality for rapid in vivo localisation and phenotyping. PLOS Neglected Tropical Diseases 12, e0006388, doi:10.1371/journal.pntd.0006388 (2018).

26 Iantorno, G. et al. The enteric nervous system in chagasic and idiopathic megacolon. Am J Surg Pathol 31, 460–468, doi:10.1097/01.pas.0000213371.79300.a8 (2007).

27 da Silveira, A. B. et al. Neuronal plasticity of the enteric nervous system is correlated with chagasic megacolon development. Parasitology 135, 1337–1342, doi:10.1017/S0031182008004770 (2008).

28 Koeberle, F. Enteromegaly and Cardiomegaly in Chagas Disease. Gut 4, 399–405, doi:10.1136/gut.4.4.399 (1963).

29 Campos, C. F. et al. Enteric Neuronal Damage, Intramuscular Denervation and Smooth Muscle Phenotype Changes as Mechanisms of Chagasic Megacolon: Evidence from a Long-Term Murine Model of *Trypanosoma cruzi* Infection. PLoS ONE 11, e0153038, doi:10.1371/journal.pone.0153038 (2016).

30 Ricci, M. F. et al. Neuronal Parasitism, Early Myenteric Neurons Depopulation and Continuous Axonal Networking Damage as Underlying Mechanisms of the Experimental Intestinal Chagas’ Disease. Front Cell Infect Microbiol 10, 583899, doi:10.3389/fcimb.2020.583899 (2020).

31 Moreira, N. M. et al. Physical exercise protects myenteric neurons and reduces parasitemia in Trypanosoma cruzi infection. Exp Parasitol 141, 68–74, doi:10.1016/j.exppara.2014.03.005 (2014).

32 Oda, J. Y. et al. Myenteric neuroprotective role of aspirin in acute and chronic experimental infections with Trypanosoma cruzi. Neurogastroenterol Motil 29, 1–13, doi:10.1111/nmo.13102 (2017).

33 de Oliveira, G. M. et al. Applicability of the use of charcoal for the evaluation of intestinal motility in a murine model of *Trypanosoma cruzi* infection. Parasitology Research 102, 747–750, doi:10.1007/s00436-007-0829-8 (2008).

34 de Souza, A. P. et al. The role of selenium in intestinal motility and morphology in a murine model of *Typanosoma cruzi* infection. Parasitology Research 106, 1293–1298, doi:10.1007/s00436-010-1794-1 (2010).

35 Messenger, L. A., Miles, M. A. & Bern, C. Between a bug and a hard place: Trypanosoma cruzi genetic diversity and the clinical outcomes of Chagas disease. Expert Rev Anti Infect Ther 13, 995–1029, doi:10.1586/14787210.2015.1056158 (2015).

36 Rassi, A. Jr., Rassi, A. & Marin-Neto, J. A. Chagas disease. Lancet 375, 1388–1402, doi:10.1016/S0140-6736(10)60061-X (2010).

37 Köberle, F. The causation and importance of nervous lesions in American trypanosomiasis. B World Health Organ 42, 739–743 (1970).

38 Boesmans, W., Lasrado, R., Vanden Berghe, P. & Pachnis, V. Heterogeneity and phenotypic plasticity of glial cells in the mammalian enteric nervous system. Glia 63, 229–241, doi:10.1002/glia.22746 (2015).

39 Hossain, E. et al. Mapping of host-parasite-microbiome interactions reveals metabolic determinants of tropism and tolerance in Chagas disease. Sci Adv 6, eaaz2015, doi:10.1126/sciadv.aaz2015 (2020).

40 Veiga-Fernandes, H. & Pachnis, V. Neuroimmune regulation during intestinal development and homeostasis. Nat Immunol 18, 116–122, doi:10.1038/ni.3634 (2017).

41 Miller, M. S., Galligan, J. J. & Burks, T. F. Accurate measurement of intestinal transit in the rat. J Pharmacol Methods 6, 211–217, doi:10.1016/0160-5402(81)90110-8 (1981).

42 Lewis, M. D. et al. Imaging the development of chronic Chagas disease after oral transmission. Sci Rep 8, 11292, doi:10.1038/s41598-018-29564-7 (2018).

43 Schmittgen, T. D. & Livak, K. J. Analyzing real-time PCR data by the comparative C(T) method. Nat Protoc 3, 1101–1108, doi:10.1038/nprot.2008.73 (2008).

